# Profiling of lysine-acetylated proteins in human urine

**DOI:** 10.1101/128207

**Authors:** Weiwei Qin, Zhenhuan Du, He Huang, Youhe Gao

## Abstract

Biomarker is the measurable change associated with a physiological or pathophysiological process, its nature is change. Contrast to the blood which is under homeostatic controls, urine reflects changes in the body earlier and more sensitive therefore is a better biomarker source. Lysine acetylation is an abundant and highly regulated post-translational modification. It plays a pivotal role in modulating diverse biological processes and is associated with various important diseases. Enrichment or visualization of proteins with specific post-translational modifications provides a method for sampling the urinary proteome and reducing sample complexity. In this study, we used anti-acetyllysine antibody-based immunoaffinity enrichment combined with high-resolution mass spectrometry to profile lysine-acetylated proteins in normal human urine. A total of 629 acetylation sites on 315 proteins were identified, including some very low-abundance proteins. This is the first proteome-wide characterization of lysine acetylation proteins in normal human urine. Our dataset provides a useful resource for the further discovery of the lysine acetylated proteins as biomarker in urine.

## 1 Introduction

Biomarker is the measurable change associated with a physiological or pathophysiological process, its nature is change(1). In contrast to the blood, which is controlled by homeostatic mechanisms, urine as waste in the body accumulates changes. We suggest it is more sensitive to detect changes in the urine may be more sensitive than in the plasma (1, 2). In addition, urine can be easily and non-invasively collected in large quantities and does not undergo significant proteolytic degradation compared to other biofluids (3). We established a urimem that adsorbs biological molecules in the urine onto a membrane (4, 5). This method makes it possible to store and archive urine samples economically on a large scale for long-term preservation. Urine may become very important for large-scale biomarker research.

Post-translational modifications (PTMs) are covalent modifications of a protein (6) and play a vital role in modulating several biological functions. In the past decade, the evolution of mass spectrometry-based proteomic methods including enrichment strategies has enabled wide-scale identification of PTMs, such as acetylation (7), glycosylation (8), and phosphorylation (9). Lysine acetylation (Kac), a dynamic and reversible PTM, was initially discovered in histones approximately 50 years ago (10). Subsequent studies were focused on chromatin remodeling for gene transcription, until the first non-histone protein, p53, was identified to be lysine-acetylated (11). In 2006, Kim et al. reported the first systematic analysis of lysine-acetylated proteins, indicating that Kac is involved in the regulation of diverse cellular pathways beyond DNA-templated processes. Subsequent studies detected abundant Kac in non-histone proteins in eukaryotes (such as in human, mouse, and *Drosophila*) (7, 12) and prokaryotes (such as *Escherichia coli, Salmonella enterica*) (13, 14).

Lysine-acetylated proteins are involved in nearly all cellular processes and are evolutionarily conserved from bacteria to mammals. These proteins are also associated with important diseases such as metabolic diseases, neurodegenerative disorders, and cardiovascular diseases(15-17).

In urine, only glycosylation and phosphorylation have been studied and applied in biomarker research(8, 18). Compared to more than 400 forms of PTMs detected in proteins(19), little is known about the PTMs of urine proteins. In this study, we used immune-affinity-based acetyllysine peptide enrichment integrated with high-resolution mass spectrometry to profile lysine-acetylated proteins in normal human urine.

## 2 Materials and methods

### 2. 1. Ethical statement

The consent procedure and the study protocol were approved by the Institutional Review Board of the Institute of Basic Medical Sciences, Chinese Academy of Medical Sciences. (Project No. 007–2014). Written informed consent was obtained from each subject prior to the study.

### 2. 2. Urine sample preparation

The first morning mid-stream urine was collected from 16 donors (8 males and 8 females), aged 24–40 years (average 30 and 27 years for males and females, respectively), routine medical examinations were normal, and females who were in menses were excluded. Next, 20 ml urine samples were filtered through a nitrocellulose membrane, urinary proteins were adsorbed onto the membrane, and the membrane was dried and stored in a vacuum bag as described previously (4). Urinary proteins were eluted from the membrane by vortexing, followed by quantification by the Bradford method for western blot analysis.

800 ml pooled urine samples (3 males and 3 females, aged 24–30 years) were centrifuged at 3500 ×*g* for 30 min. After discarding the debris, the supernatant was centrifuged at 12,000 ×*g* for 30 min. After the removal of precipitates, urinary proteins were fractionated by acetone and re-dissolved in lysis buffer (7 M urea, 2 M thiourea, 120 mM dithiothreitol, and 40 mM Tris), and then quantified by the Bradford method for immune-affinity enrichment and LC-MS/MS analysis.

### 2. 3. Western blot analysis

Urinary proteins from each sample (30 μg, n = 16) were loaded on 12% SDS-PAGE and transferred to a polyvinylidene difluoride (PVDF) membrane. After blocking in TBST buffer (1X TBS containing 0. 1% Tween 20) with 5% (w/v) nonfat milk for 40 min at room temperature, the membranes were incubated with diluted primary antibodies (dilution 1:1000) (PTM Biolabs, Chicago, IL, USA) at 4°C with gentle shaking overnight. Next, the membrane was probed with secondary antibodies (dilution 1:5000) coupled to horseradish peroxidase. Signals were revealed by enhanced chemiluminescence, and the results were scanned and analyzed using an Image Quant 400TM Imager (GE Healthcare, Little Chalfont, UK). Finally, the membrane was stained with Coomassie Brilliant Blue.

### 2. 4. Protein digestion

5 mg urinary proteins were used for digestion, the protein solution was reduced with 10 mM DTT for 1 h at 37°C and alkylated with 20 mM IAA for 45 min at room temperature in the dark. For trypsin digestion, the protein sample was diluted by adding 100 mM ammonium bicarbonate to make the urea concentration less than 2 M. Finally, trypsin was added at a 1:50 trypsin-to-protein mass ratio for the first digestion overnight and 1:100 trypsin-to-protein mass ratio for the second 4-h digestion.

### 2. 5. Off-line high-pH HPLC separation

2 mg tryptic peptides was then fractionated by high pH reverse-phase HPLC using Agilent 300Extend C18 column (5 μm particles, 4.6 mm ID, 250 mm length, Santa Clara, CA, USA). Briefly, peptides were first separated over a gradient of 2–60% acetonitrile in 10 mM ammonium bicarbonate, pH 10.0 for 80 min into 80 fractions. Next, the peptides were combined into 7 fractions and dried by vacuum centrifugation.

### 2. 6. Enrichment of lysine-acetylated peptides

To enrich Kac peptides, tryptic peptides dissolved in NETN buffer (100 mM NaCl, 1 mM EDTA, 50 mM Tris-HCl, 0.5% NP-40, pH 8.0) were incubated with pre-washed antibody beads (PTM Biolabs) at 4°C overnight with gentle shaking. The beads were washed four times with NETN buffer and twice with ddH_2_O. The bound peptides were eluted from the beads with 0.1% TFA. The eluted fractions were combined and vacuum-dried. The resulting peptides were cleaned with C18 ZipTips (Millipore, Billerica, MA, USA) according to the manufacturer’s instructions, followed by LC-MS/MS analysis.

### 2. 7. LC-MS/MS analysis

The lyophilized peptides were re-suspended in 0.1% formic acid and subjected to chromatography using an EASY-nLC 1000 UPLC system (Thermo Scientific, Waltham, MA, USA). The peptides were separated on a reversed-phase analytical column (Acclaim PepMap RSLC, Thermo Scientific). Elution was performed over a gradient of 7–35% buffer B (0.1% formic acid in 98% ACN; flow rate, 0.3 μL/min) for 40 min. A Q Exactive^TM^ Plus mass spectrometer (Thermo Scientific) was used to analyze the fractions. The MS data were acquired using the high-sensitivity mode with the following parameters: 20 data-dependent MS/MS scans per full scan, full scans acquired at a resolution of 70,000 and MS/MS scans at a resolution of 17,500, dynamic exclusion (exclusion duration 30 s), MS scans over a range of 350–1800 m/z.

### 2. 8. Data processing

The MS raw data were processed using the PEAKS Studio (version 7.5, Bioinformatics Solution Inc., Waterloo, Canada). High-quality de novo sequence tags were then used by PEAKS DB(20) to search the human proteome database (UniProtKB/Swiss-Prot release 2014-01-10). The fasta file contained 20120 protein sequences. A mass tolerance of 10 ppm and 0.02 Da was set for the precursor ions and fragment ions, respectively. Cysteine carbamidomethylation was set as fixed modification, oxidation (M), protein N-terminal and lysine acetylation as variable modifications. Trypsin allowing 3 missed cleavages was chosen as the enzyme. A decoy database was also searched to calculate the false-discovery rate (FDR) using the decoy-fusion method. Result filtration parameters: peptide-spectrum matches FDR < 0.1 %; Peptide −10 lgP ≥ 27.5 (FDR 0.2%); Protein –10 lgP ≥ 28 (FDR 0.0%), and proteins unique peptides ≥ 1; AScore ≥ 59 considers as confident Kac site.

### 2. 9. Motif and secondary structure analysis

Software motif-x was used to analyze the model of sequences with amino acids in specific positions of acetylated lysine-13-mers (6 amino acids upstream and downstream of the acetylation site) in all protein sequences (21). The IPI human proteome was used as an extension database, while other parameters were set to default values. The local secondary structures of lysine-acetylated proteins were predicted by NetSurfP (22).

### 2. 10. Protein networks and functional analysis

All identified lysine-acetylated urinary proteins were subjected to network and functional analyses using ingenuity pathway analysis (IPA) version 9.0 (http://www.ingenuity.com).

## 3 Results and discussion

### 3. 1 Western blot analysis of lysine-acetylated urinary proteins of healthy humans

In order to explore the status of lysine acetylation in the urine proteome of healthy humans, western blot analysis was performed using a pan anti-acetyllysine antibody. The results showed that multiple protein bands with a wide range of molecular weights were detected. In addition to histones (11–15 kDa and 26 kDa), many non-histone proteins with clear bands were observed (Fig 1B and Fig. S1), indicating that lysine acetylation is highly abundant in human urine. Furthermore, the acetylated proteins showed different patterns among different healthy individuals. This suggests that lysine acetylation levels reflect various statuses of healthy people.

**Figure 1.**
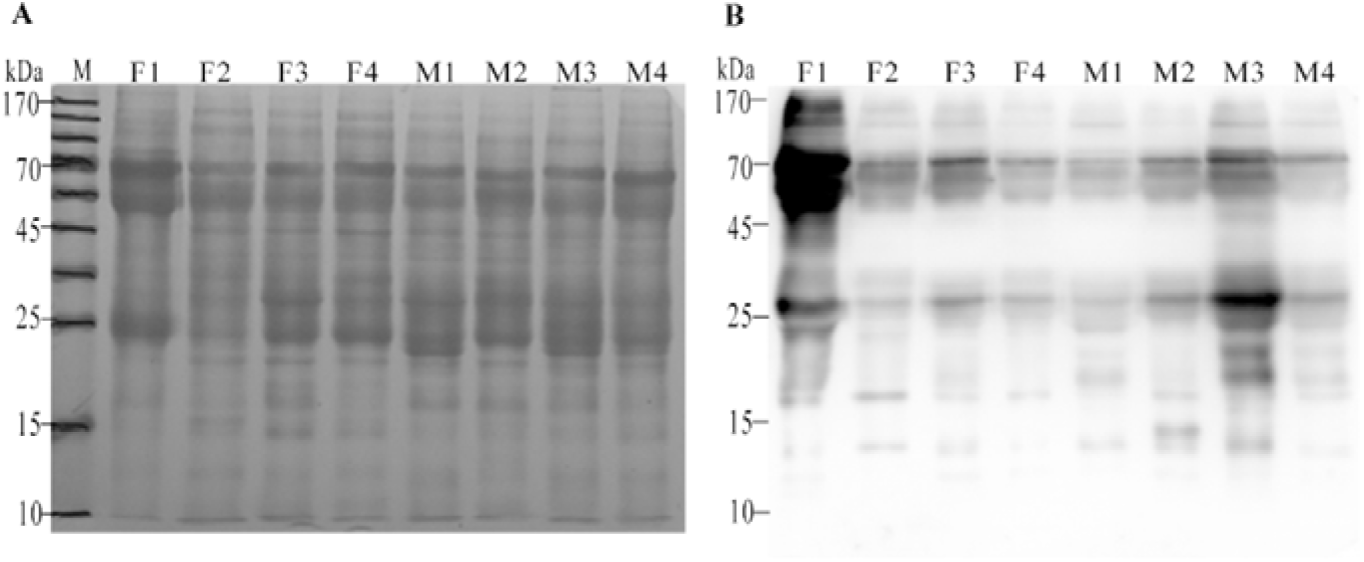
Overview of lysine acetylation in urine proteome of healthy humans. (A) SDS-PAGE analysis of urinary proteins followed by transfer to PVDF membrane and staining with Coomassie Brilliant Blue; (B) Western blotting analysis of urinary proteins. (M: male, F: female).

### 3. 2. Profiling of lysine-acetylated urinary proteins

Because lysine acetylation plays a vital role in modulating various molecular processes and widely exists in the urine, we comprehensively profiled lysine-acetylated proteins in the urine using immune-affinity-based acetyllysine peptide enrichment integrated with LC-MS/MS.

We first identified 761 acetylated peptides (Peptide-Spectrum Matches FDR < 0.1 %) (Table S1) matched on 315 lysine-acetylated proteins (Unique peptides ≥ 1) with 629 Kac sites (AScore ≥ 59) in normal human urine (Table S2). Among these proteins, 18 contained five or more Kac sites, accounting for 5.7% of the total detected Kac proteins with an average degree of acetylation of 2.

When these 315 proteins were searched in the most comprehensive urinary database (www.urimarker.com/urine/) (Fig 2A), 242 were found in 1D group (proteins detectable when separated by 1D LC) and 71 in the 2D and 3D group (2D: high-pH RPLC coupled LC- MS/MS, 3D: GELFrEE/LP-IEF fraction prior to 2D), which were less abundant proteins; 2 were not found in this database.

**Figure 2.**
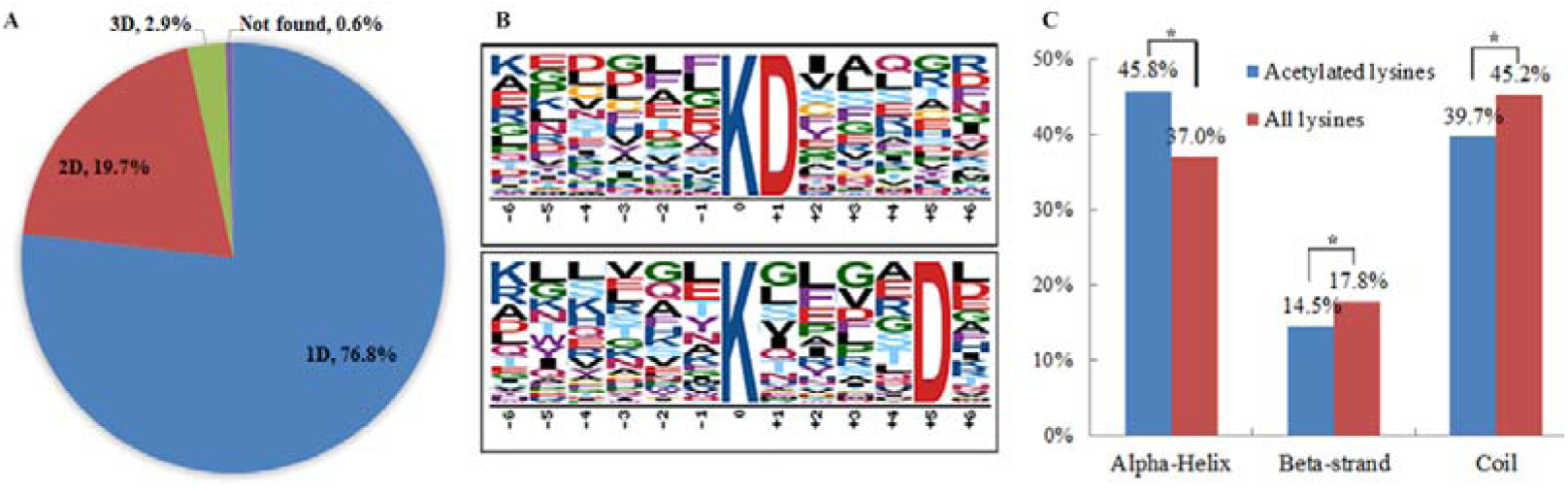
Statistics of the lysine-acetylated proteins. (A) Estimation of relative abundance of urinary lysine-acetylated proteins. When the 315 proteins were searched in the most comprehensive urinary database, 242 were found in the 1D group (proteins detectable when separated by 1D LC) and 71 in the 2D and 3D group (2D: high-pH RPLC coupled LC- MS/MS, 3D: GELFrEE/LP-IEF fraction prior to 2D), which were less abundant proteins. (B) Acetylation motifs and conservation of acetylation sites. (C) Distribution of acetylated and non-acetylated lysine residues in protein secondary structures (*p < 0.01).

This is the first proteome-wide characterization of lysine acetylation proteins in normal human urine. A total of 629 acetylation sites on 315 proteins were identified, including some very low-abundance proteins. Enrichment of acetylated peptides and a high pH pre-fraction of unenriched peptides vastly reduced the sample complexity and increased the dynamic range. For albuminuria occurring in some kidney diseases, the masking effects of highly abundant proteins and ion suppression were much stronger. To address this issue, a combination of two types of PTMs, such as *N*-glycosylated proteins and lysine-acetylated peptide enrichment, may be used.

### 3. 3. Analysis of lysine acetylation sites

To examine the relative abundance of the amino acids flanking the acetylation sites, we analyzed the 761 lysine-acetylated peptides using Motif-X software. A negatively charged residue, aspartate, at the +1 and +5 positions of acetyllysine was identified (Fig 2B). According to previous studies of human lysine acetylation (23), aspartate was found to be enriched at the -1, -2, and +1 positions of the lysine acetylation site. To further explore the relationship between the lysine acetylation and protein secondary structures, we performed structural analysis of all acetylated proteins using NetSurfP. Acetylated lysine was found in alpha-helices approximately 8.8% more frequently than the average lysine and in the coil approximately 5.5% less frequently (p < 0.01) (Fig 2C). Thus, acetylated lysine appears to be enriched in structured regions and depleted in unstructured regions, which is in concordance with previous reports (7, 12).

### 3. 4. Functional analysis of lysine-acetylated proteins in urine

Classification analysis of the lysine-acetylated proteins identified in urine was performed using the IPA tool (http://www.ingenuity.com). As shown in Fig 3A, 42.4% of lysine-acetylated proteins were annotated in the cytoplasm, 33.3% in the extracellular space, 18.8% in the plasma membrane, and 5.2% in the nucleus. The proportion of cytoplasm proteins in the lysine acetylome was much higher than that in the whole urinary proteome (17%) (24), while the other proportions of extracellular space or plasma membrane proteins were much lower than that in the whole urinary proteome (38% and 31%) (24). The major types of lysine-acetylated proteins were enzymes (enzyme 25.9%, peptidase 11%, kinase 2.3%, phosphatase 1.9%, and transporters 10%) (Fig 3B).

**Figure 3.**
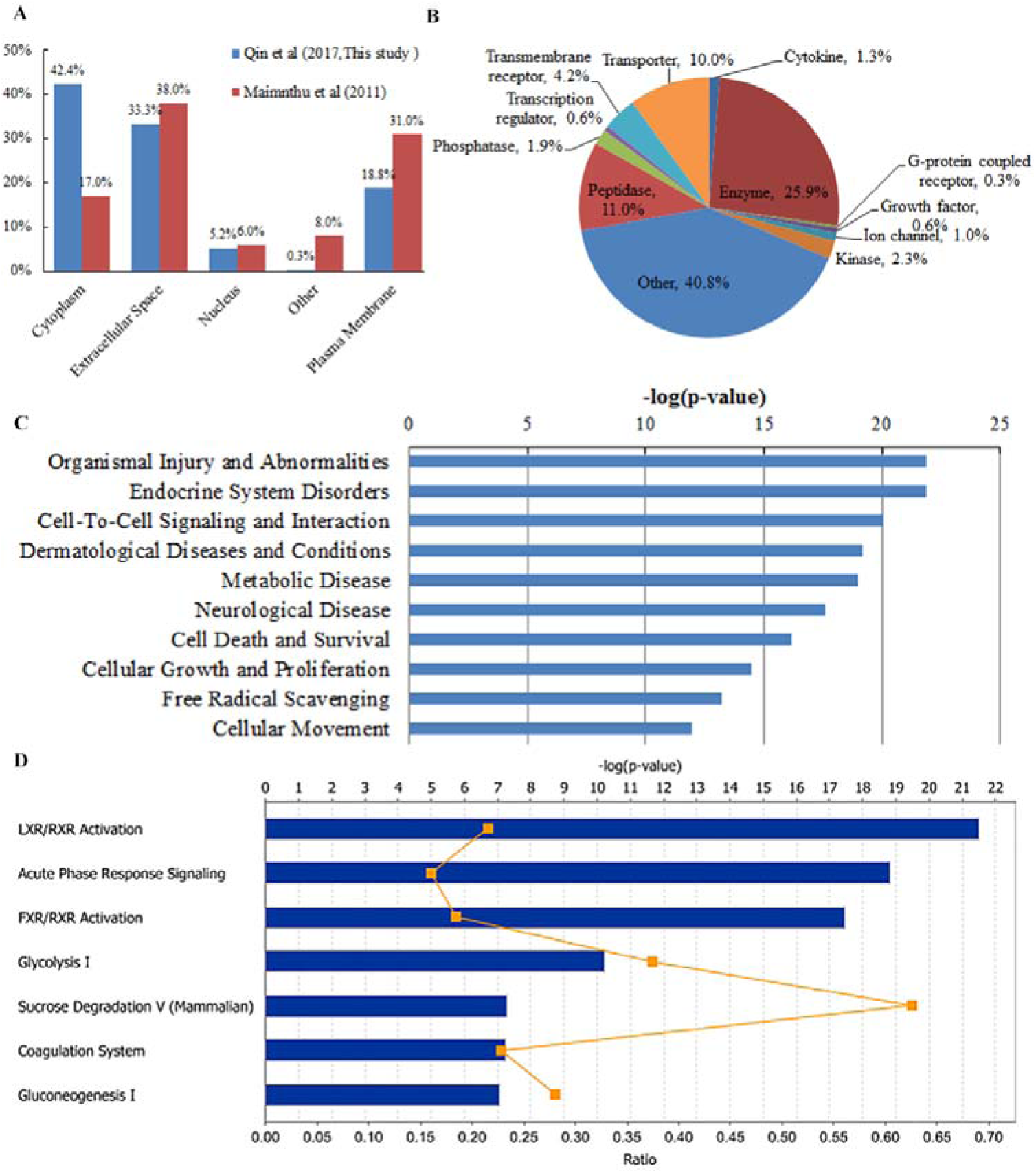
Functional analysis of the lysine acetylome in urine. The lysine-acetylated proteins identified in this study were classified according to their (A) subcellular location, (B) protein family, (C) enriched bio functions and diseases, and (D) canonical pathways and networks.

Enriched bio functions and diseases were analyzed using the IPA tool. As shown in Fig 3C, the notable bio functions of these proteins were cell-to-cell signaling and interaction (87 proteins), cell death and survival (118 proteins), cellular growth and proliferation (115 proteins), free radical scavenging (28 proteins), and cellular movement (76 proteins). The major diseases and disorders associated with these proteins were found to be endocrine system disorders (110 proteins), organismal injury and abnormalities (299 proteins), dermatological diseases and conditions (73 proteins), metabolic disease (95 proteins), and neurological diseases (116 proteins). These results provide valuable information regarding lysine acetylation in the urine and may be useful in biomarker studies these diseases and disorders.

The main canonical pathways (Fig 3D) in which the lysine-acetylated proteins participated, including LXR/RXR activation, acute phase response signaling, and FXR/RXR activation, showed the same rankings in the whole urinary proteome. Among these top 7 networks, three were associated with glucose metabolism, such as glycolysis I, sucrose degradation V (mammalian), and gluconeogenesis I.

Lysine acetylated proteins in the urine may reflect protein acetylation status in cells. Discarding acetylated proteins may provide a way to modulate levels of lysine acetylated proteins in cells. Our dataset provides a useful resource for the further discovery of the lysine acetylated proteins as biomarker in urine.

## Acknowledgements

Funding: This work was supported by the National Key Research and Development Program of China (grant number 2016YFC1306300); the National Basic Research Program of China (grant number 2013CB530850); and funds from Beijing Normal University (grant numbers 11100704, 10300-310421102).

## Supplementary Material

The flowing supporting information and the raw MS/MS data have been uploaded to the data repository. (https://figshare.com/s/4983659b9db1d69b7b8c)

Supporting Fig S1. Overview of lysine acetylation in urine proteome of healthy humans. (A) SDS-PAGE analysis of urinary proteins followed by transfer to PVDF membrane and staining with Coomassie Brilliant Blue; (B) Western blotting analysis of urinary proteins. (M: male, F: female).

Supporting Table S1. Acetylated peptides (Peptide-Spectrum Matches FDR < 0.1 %).

Supporting Table S2. Acetylated proteins (Unique peptides ≥ 1) with 629 Kac sites (AScore ≥ 59) in normal human urine.

Supporting Spectra S1. The represent MS/MS spectra for 761 lysine acetylated peptides.

Supporting Spectra S2. The remaining MS/MS spectra for 761 lysine acetylated peptides.

## References

1. Gao Y. Urine-an untapped goldmine for biomarker discovery? Sci China Life Sci. 2013; 56(12): 1145–6.

2. Li M, Zhao M, Gao Y. Changes of proteins induced by anticoagulants can be more sensitively detected in urine than in plasma. Sci China Life Sci. 2014; 57: 649–54.

3. Ste´phane D, Anne P, Benjamin B, Harald M. Urine in Clinical Proteomics. Mol Cell Proteomics. 2008; 7: 1850–62.

4. Jia L, Liu X, L L, al e. Urimem, a membrane that can store urinary proteins simply and economically, makes the large-scale storage of clinical samples possible. Sci China Life Sci. 2014; 57: 336–9.

5. Zhang F, Cheng X, Yuan Y, Wu J, Gao Y. Urinary microRNA can be concentrated, dried on membranes and stored at room temperature in vacuum bags. PeerJ. 2015; 3: e1082.

6. Mann M, Jensen ON. Proteomic analysis of post-translational modifications. Nature Biotechnology 2003; 21: 7.

7. Kim SC, Sprung R, Chen Y, Xu Y, Ball H, Pei J, et al. Substrate and functional diversity of lysine acetylation revealed by a proteomics survey. Molecular cell. 2006; 23(4): 607–18.

8. Wang L, Li F, Sun W, Wu S, Wang X, Zhang L, et al. Concanavalin A-captured glycoproteins in healthy human urine. Molecular & cellular proteomics: MCP. 2006; 5(3): 560–2.

9. Olsen JV, Blagoev B, Gnad F, Macek B, Kumar C, Mortensen P, et al. Global, in vivo, and site-specific phosphorylation dynamics in signaling networks. Cell. 2006; 127(3): 635–48.

10. Vidali G, Gershey EL, Allfrey VG. Chemical Studies of Histone Acetylation. The Distribution of ɛN-acetyllysine in Calf Thymus Histones. J Biol Chem. 1968; 243: 6361–6.

11. Gu W, Roeder GR. Activation of p53 Sequence-Specific DNA Binding by Acetylation of the p53 C-Terminal Domain. Cellular signalling. 1997; 90: 595–606.

12. Choudhary C, Kumar C, Gnad F, Nielsen ML, Rehman M, Walther TC, et al. Lysine acetylation targets protein complexes and co-regulates major cellular functions. Science. 2009;325(5942):834–40.

13. Zhang J, Sprung R, Pei J, Tan X, Kim S, Zhu H, et al. Lysine acetylation is a highly abundant and evolutionarily conserved modification in Escherichia coli. Molecular & cellular proteomics: MCP. 2009; 8(2): 215–25.

14. Wang Q, Zhang Y, Yang C, Xiong H, Lin Y, Yao J, et al. Acetylation of metabolic enzymes coordinates carbon source utilization and metabolic flux. Science. 2010;327(5968):1004–7.

15. Menzies KJ, Zhang H, Katsyuba E, Auwerx J. Protein acetylation in metabolism - metabolites and cofactors. Nature reviews Endocrinology. 2016; 12(1): 43–60.

16. Voelter-Mahlknecht S. Epigenetic associations in relation to cardiovascular prevention and therapeutics. Clinical epigenetics. 2016; 8: 4.

17. Pons D, de Vries FR, van den Elsen PJ, Heijmans BT, Quax PH, Jukema JW. Epigenetic histone acetylation modifiers in vascular remodelling: new targets for therapy in cardiovascular disease. European heart journal. 2009; 30(3): 266–77.

18. Li QR, Fan KX, Li RX, Dai J, Wu CC, Zhao SL, et al. A comprehensive and non-prefractionation on the protein level approach for the human urinary proteome: touching phosphorylation in urine. Rapid communications in mass spectrometry: RCM. 2010; 24(6): 823–32.

19. Rotilio D, Della Corte A, D’Imperio M, Coletta W, Marcone S, Silvestri C, et al. Proteomics: bases for protein complexity understanding. Thrombosis research. 2012; 129(3): 257–62.

20. Zhang J, Xin L, Shan B, Chen W, Xie M, Yuen D, et al. PEAKS DB: de novo sequencing assisted database search for sensitive and accurate peptide identification. Molecular & cellular proteomics: MCP. 2012;11(4):M111 010587.

21. Chou MF, Schwartz D. Biological sequence motif discovery using motif-x. Curr Protoc Bioinformatics. 2011;Chapter 13:Unit 13 5–24.

22. Petersen B, Petersen TN, Andersen P, Nielsen M, Lundegaard C. A generic method for assignment of reliability scores applied to solvent accessibility predictions. BMC structural biology. 2009; 9: 51.

23. Zhu X, Liu X, Cheng Z, Zhu J, Xu L, Wang F, et al. Quantitative Analysis of Global Proteome and Lysine Acetylome Reveal the Differential Impacts of VPA and SAHA on HL60 Cells. Scientific reports. 2016; 6: 19926.

24. Marimuthu A, O’Meally RN, Chaerkady R, Subbannayya Y, Nanjappa V, Kumar P, et al. A comprehensive map of the human urinary proteome. Journal of proteome research. 2011; 10(6): 2734–43.

